# TARO: tree-aggregated factor regression for microbiome data integration

**DOI:** 10.1101/2023.10.17.562792

**Authors:** Aditya K. Mishra, Iqbal Mahmud, Philip L. Lorenzi, Robert R. Jenq, Jennifer A. Wargo, Nadim J. Ajami, Christine B. Peterson

**Affiliations:** Department of Genomic Medicine, The University of Texas MD Anderson Cancer Center, Houston, TX 77030, United States; Platform for Innovative Microbiome and Translational Research (PRIME-TR), The University of Texas MD Anderson Cancer Center, Houston, TX 77030, United States; Department of Bioinformatics and Computational Biology, The University of Texas MD Anderson Cancer Center, Houston, TX 77030, United States; Department of Surgical Oncology, The University of Texas MD Anderson Cancer Center, Houston, TX 77030, United States; Department of Biostatistics, The University of Texas MD Anderson Cancer Center, Houston, TX 77030, United States

**Keywords:** factor model, reduced-rank regression, microbiome data, data integration, feature aggregation

## Abstract

**Motivation:** Although the human microbiome plays a key role in health and disease, the biological mechanisms underlying the interaction between the microbiome and its host are incompletely understood. Integration with other molecular profiling data offers an opportunity to characterize the role of the microbiome and elucidate therapeutic targets. However, this remains challenging to the high dimensionality, compositionality, and rare features found in microbiome profiling data. These challenges necessitate the use of methods that can achieve structured sparsity in learning cross-platform association patterns.

**Results:** We propose Tree-Aggregated factor RegressiOn (TARO) for the integration of microbiome and metabolomic data. We leverage information on the phylogenetic tree structure to flexibly aggregate rare features. We demonstrate through simulation studies that TARO accurately recovers a low-rank coefficient matrix and identifies relevant features. We applied TARO to microbiome and metabolomic profiles gathered from subjects being screened for colorectal cancer to understand how gut microrganisms shape intestinal metabolite abundances.

**Availability and implementation:** The R package `TARO` implementing the proposed methods is available online at https://github.com/amishra-stats/taro-package.

## Introduction

The human microbiome consists of a diverse community of microorganisms, including bacteria, fungi, and viruses, that populate various sites in the body. The microbiome plays a key role in many normal biological processes in the host including digestion and immune regulation, while dysbiosis, or disruption of healthy microbiome composition, has been linked to disease risk across a range of conditions, including heart disease, diabetes, and cancer [Hou et al., 2022]. Although mechanisms of host-microbiome interaction remain incompletely understood, one avenue for the influence of the microbiome on host disease processes is through the production of metabolites [Cullin et al., 2021]. Characterizing the influence of the microbiome on the metabolome requires integrative analysis of these high-throughput data types. However, this task is challenging for several reasons: both microbiome and metabolite data are high-dimensional, with thousands of features measured in each sample; microbiome data are compositional, which means that each sample has a fixed sum constraint; and microbiome data are zero-inflated, which means that a feature observed in one sample is often not observed in other samples, resulting in rare features with a large number of observed zero values.

Various methods have been proposed for cross-platform integration of microbiome and metabolomics data including correlation and network inference approaches. A naïve method popular in practice is to test for pairwise associations between individual features using Pearson’s or Spearman’s correlation; however, this approach creates a high multiple testing burden. Classical multivariate methods, such as canonical correlation analysis (CCA) and co-inertia analysis (CIA), are attractive alternatives but require that the number of samples *n* is larger than the number of variables. Given the high dimensionality of data sets in the high-throughput era, sparse versions of CCA and CIA have been developed to resolve this limitation [Witten and Tibshirani, 2009, Min et al., 2019]. Network inference methods based on the graphical modeling framework have also been proposed as an approach for the integration of microbiome data with high-dimensional covariates [Yang et al., 2017, Osborne et al., 2022]; an advantage of these methods is that they aim to capture direct associations by focusing on conditional, rather than marginal, correlations. However, none of these methods directly handle the challenge of rare features.

Here, we frame the challenge of integrating microbiome and metabolite data as a factor regression model, with the microbiome profiles as the predictor and metabolite profiles as the response. We propose Tree-Aggregated factor RegressiOn (TARO), building on the reduced-rank regression framework to enable the discovery of interpretable latent factors with flexible aggregation of rare features. Our proposed approach leverages information on the phylogenetic tree to enable aggregation of features in a data-adaptive manner, collapsing rare features into aggregated features that are less zero-inflated. Our proposed method offers a comprehensive solution to the challenges of high-dimensionality, compositionality, and rare features. In Section 2, we provide a description of the proposed model and estimation procedure. In Section 3, we compare the performance of TARO to alternative methods through simulation studies and apply TARO to integrate microbiome and metabolomics data from a real-world study on colorectal cancer [Yachida et al., 2019]. Finally, we conclude with a discussion in Section 4.

## Materials and methods

### TARO model

Our proposed method builds on the multivariate regression framework to relate the microbiome and metabolomic profiling data. We assume that the observed microbiome data consists of abundances for *p* features across *n* samples. The features may correspond to taxonomic units quantified through marker gene sequencing, such as amplicon sequence variants (ASVs) or operational taxonomic units, or more generally to any functional or taxonomic read-outs. We denote the microbial abundance table as **W** = [*w*_*ij*_]_*n×p*_ = [**w**_1_, …, **w**_*n*_]^T^. Importantly, due to the methods employed for the generation and processing of the genomic sequences, the observed data are compositional; this means that the observed counts can only be interpreted on a relative scale [Gloor et al., 2017]. Regression models with microbiome features as the predictor typically rely on data transformations to address this challenge [Aitchison and Bacon-Shone, 1984]. Here, we first apply total sum scaling (TSS), which entails dividing each count *w*_*ij*_ by the total number of counts for its sample ∑_*j*_ *w*_*ij*_. Recent work has shown that TSS scaling, although quite simple, tends to perform better in practice than other normalization methods [Mallick et al., 2021]. This results in a relative abundance matrix 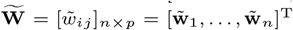 such that 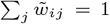 for *i* = 1, …, *n*. We then apply a log transform to obtain the matrix **X** = [*x*_*ij*_]_*n×p*_ where 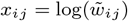. To avoid numerical issues with exact zeros, we add a pseudocount of 1 to the count matrix **W** prior to scaling. Importantly, the resulting *p* features are not independent, as there are only *p −*1 degrees of freedom due to the original sum constraint.

We now consider the formulation of the regression model relating the microbial profiling data **X** to the metabolite abundances. We let **Y** = [*y*_*ik*_]_*n×q*_ = [**y**_1_, …, **y**_*n*_]^T^ *∈* ℝ^*n×q*^ represent the metabolite abundances; since metabolomic data are often highly skewed, the *y*_*ik*_ may be taken as the log-transformed concentration values. The metabolite abundances can be modeled as a function of a set of covariates **Z**_*n×m*_ and the microbiome profiles **X**_*n×p*_ via the multivariate regression:

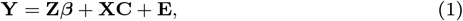

where ***β***_*m×q*_ represents the matrix of effects of the clinical covariates on the metabolites, **C**_*p×q*_ is the matrix of effects of the microbiome features on the metabolites, and **E**_*n×q*_ = [*e*_*ik*_]_*n×q*_ is the error matrix. We include an intercept in the model by setting the first column of **Z** to be **1**_*n*_. The remaining columns correspond to clinical variables we wish to include as adjusters in the model and are not subject to selection. We therefore do not impose any regularization on ***β***.

The novelty of the TARO method lies in how we estimate the coefficient matrix **C** to handle the compositionality of the microbiome profiles, aggregate rare features, and achieve sparsity. Due to the fixed sum constraint within each sample, the *p* microbiome predictor variables are not independent; this means that an additional constraint is needed to ensure identifiability of the coefficient matrix **C**. Following Lin et al. [2014], we incorporate a zero-sum constraint on each row of **C**:

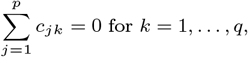

which can be written in matrix form as 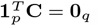.

Next, we consider how to aggregate rare features; this is a critical challenge in microbiome data analysis since the observed fine-resolution microbiome features typically include features that are non-zero in only a few samples. Many existing approaches for microbiome regression collapse the observed features to a higher taxonomic rank, typically the genus level [Lin et al., 2014, Liu et al., 2022]. For example, the counts for ASV1 and ASV2 could be summed to obtain the abundance for their parent genus in the taxonomic tree *𝒯*. This can be carried out to obtain abundances for any internal node in the tree. Suppose we obtain a new aggregated feature for the *i*th subject 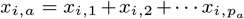, where *p*_*a*_ denotes the number of leaf nodes descending from the parent node *a*. As noted in Yan and Bien [2021], 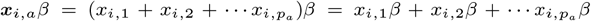. Effectively, this means that learning a model where some features have exactly equal coefficients *β* corresponds to aggregating the original features into less zero-inflated groupings. Here, we build on the work of Yan and Bien [2021] and Bien et al. [2021] to allow flexible estimation of the microbiome coefficients, to allow grouping of rare features when the data supports their having equivalent effects on the outcome.

Following Yan and Bien [2021], we denote the internal nodes of the taxonomic tree 𝒯 using an index set *u ∈ {*1, …, |*𝒯* | *−* 1*}*, excluding the root node, and let **A**_*p×*(|*𝒯* |*−*1)_ = [*a*_*ju*_] indicate the ancestry of each observed feature, where the entry *a*_*ju*_ = 1 if microbiome feature *j* belongs to the set of leaves descending from node *u*, and *a*_*ju*_ = 0 otherwise. To enable flexible feature aggregation, we rewrite the coefficient matrix **C** as follows:

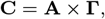

where **Γ**_(|*𝒯*|*−*1)*×q*_. This reparameterization results aggregated features as 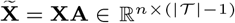. We can then expand the model (1) as:

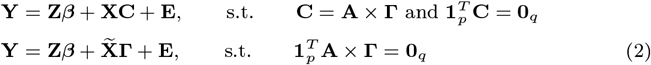

Since **Γ**_(|*𝒯* |*−*1)*×q*_ is high-dimensional, it is critical to leverage assumptions on its structure to reduce the number of parameters to be estimated. To do so, we build on the framework of reduced-rank regression [Izenman, 1975, Chen and Huang, 2012]. The key idea of reduced-rank regression is to impose a constraint on the rank, or number of linearly independent rows or columns, of **Γ**, such that *r* = rank(**Γ**) *<* min((|*𝒯* | *−* 1), *q*). Following recent advances in factor regression modeling [Mishra et al., 2017], we express **Γ** as a low-rank and sparse coefficient matrix using the components from the singular value decomposition (SVD):

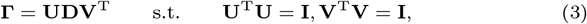

where the left singular vectors are given by **U** = [**u**_1_, …, **u**_*r*_], the right singular vectors are given by **V** = [**v**_1_, …, **v**_*r*_] and the singular values are given by diag(**D**) = [*d*_1_, …, *d*_*r*_]. A schematic overview of the TARO model is shown in Figure 1.

**Fig. 1:**
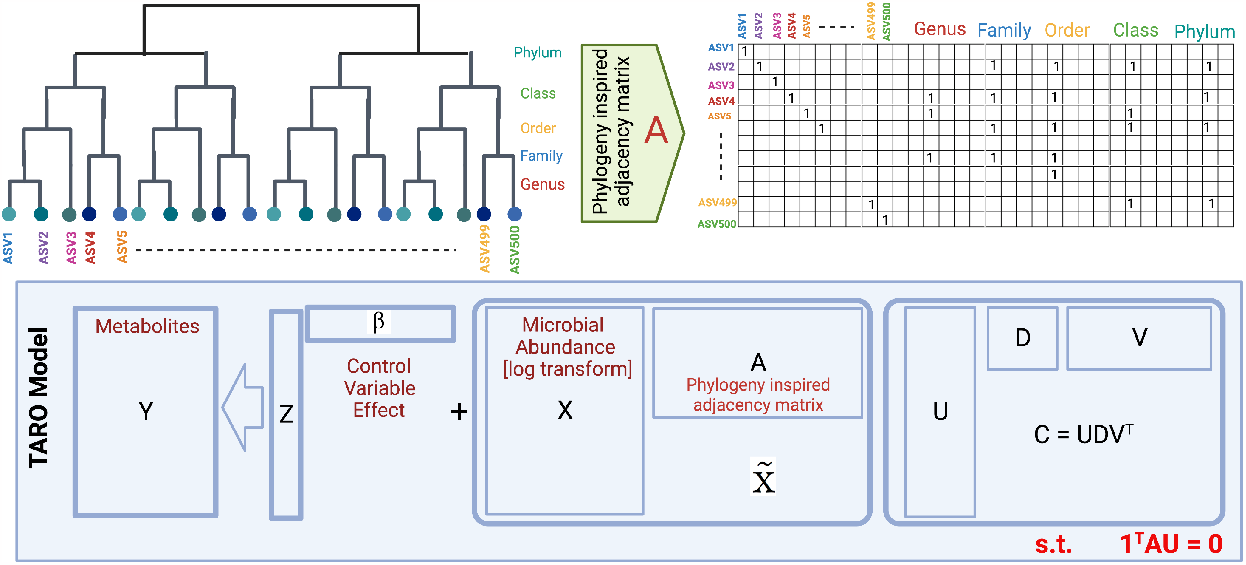
Overview of the TARO model.

### Sequential estimation of TARO

We now describe our efficient computational procedure for obtaining estimates of the model parameters. With the rank *r* of the coefficient matrix **Γ** specified, the model parameters can be estimated by solving the optimization problem:

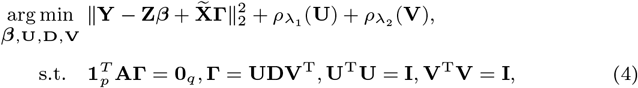

where 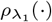 and 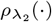 are sparsity inducing penalties with tuning parameters *λ*_1_ and *λ*_2_, respectively. In high-dimensional settings, sparse estimates of the singular vectors facilitate better model interpretation. With the rank of **C** unknown and an orthogonality constraint on the singular vectors *{***U, V***}*, joint estimation of the parameters is a notoriously intractable problem [Chen, 2011, Mishra et al., 2017, 2021]. However, when orthogonality constraints are dropped, the singular vectors become unidentifiable. As a result, the sparsity pattern in the singular vectors is not unique, which hinders model interpretation. Following the work of Mishra et al. [2017], we overcome the challenge by using a sequential approach to estimate the model parameters. Under this approach, we express **Γ** as the sum of *r* unit-rank matrices:

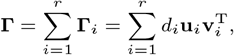

where *{d*_*i*_, **u**_*i*_, **v**_*i*_*}* are SVD components. The estimation procedure then estimates the SVD components *{d*_*i*_, **u**_*i*_, **v**_*i*_*}* of **Γ** in sequential order.

#### Step 1

Extract the first components: With the aim to estimate *{d*_1_, **u**_1_, **v**_1_, ***β****}*, we solve the optimization problem:

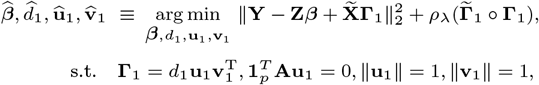

where 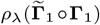 is a weighted adaptive elastic-net penalty [Mishra et al., 2017] with weights 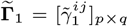 inducing sparsity of both the left and right singular vectors *{***u**_1_, **v**_1_*}*. Details on the construction of the weights 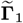 and the formulation of the weighted penalty are given in Supplementary Material Section S1.

#### Step k

Extract the *k*th components: With the aim to estimate *{d*_*k*_, **u**_*k*_, **v**_*k*_, ***β****}*, we solve the optimization problem:

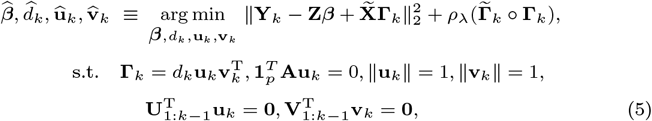

where 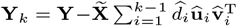 is the deflated response matrix and 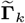 is the weight matrix for constructing the sparsity inducing penalty. Motivated by the constraints in the optimization problem (4), the additional constraints 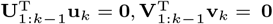 are required for imposing orthogonality on the estimate of the singular vectors. Such constraints are necessary in the optimization (4) for the estimates to be identifiable. However, in the sequential approach one can safely drop the additional constraints and still have an estimate of the singular vectors with a unique sparsity pattern. Hence, to extract the *k*th SVD components, we solve the optimization problem:

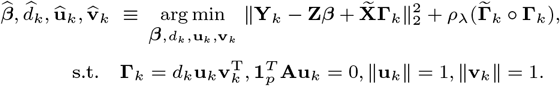

We write the general form of the optimization problem in any *k*th step of the sequential procedure as:

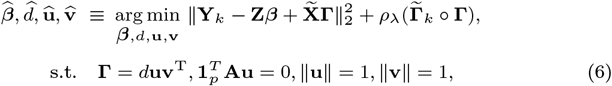

where 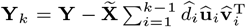 is the deflated response matrix. We conveniently represent the unit-rank estimation problem for TARO as URE-TARO(*d*, **u, v**, 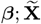, **Z, Y**_*k*_, **A**, 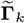).We summarize the sequential procedure for parameter estimation in Algorithm 1, with additional details provided in Section S2 of the Supplementary Material.

##### Algorithm 1 Tree-Aggregated factor RegressiOn (TARO)

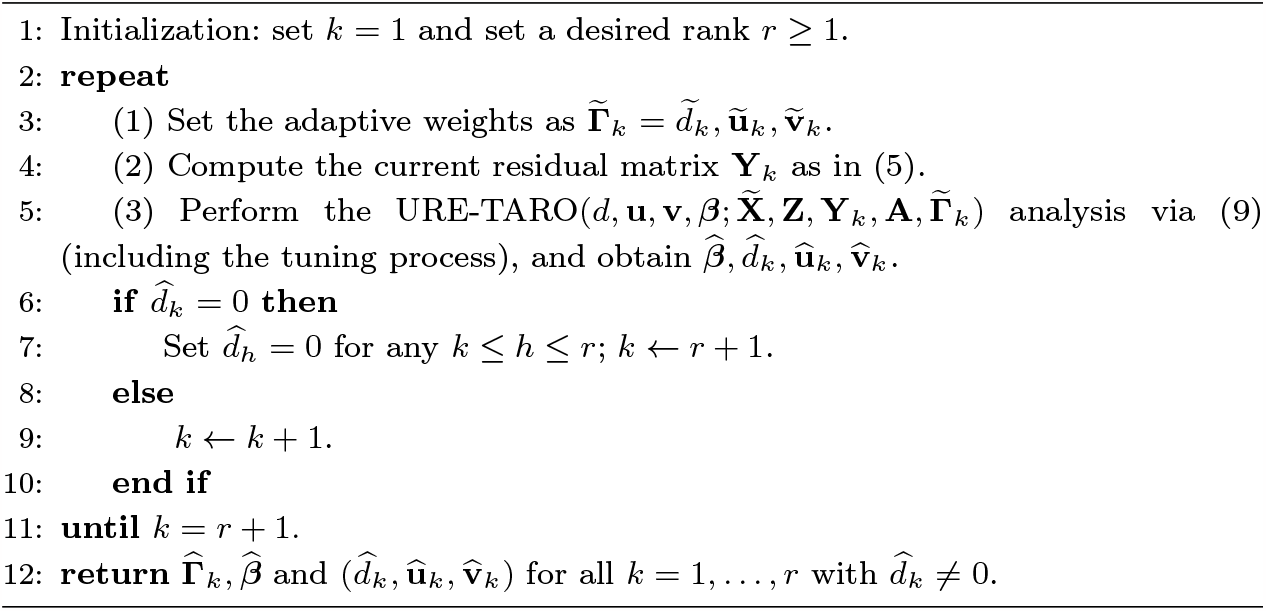

## Results

### Simulation study

To assess the performance of TARO in comparison to alternative approaches, we carried out a series of simulation studies. The generation of synthetic microbiome profiles is a challenging task due to the complex data structure of microbiome compositional profiles obtained from specimens. To simulate realistic microbial abundances, we relied on SparseDOSSA2 [Ma et al., 2021], which utilizes a real data template as a target for the marginal feature distributions. As our template, we relied on the stool profiles from the expanded Human Microbiome Project [Lloyd-Price et al., 2017]. We then scaled and log-transformed the resulting counts **W** to obtain our **X** matrix. We generated a coefficient matrix **C** with true rank *r* = 3 based on the unit-rank components *{d*_*i*_, **u**_*i*_, **v**_*i*_*}* for 1 *≤ i ≤ r*. We set *d*_1_ = 4, *d*_2_ = 3, and *d*_3_ = 2, and simulated sparse and nearly orthogonal **U** = [**u**_1_, **u**_2_, **u**_3_] and **V** = [**v**_1_, **v**_2_, **v**_3_] matrices. We utilized a taxonomic tree 𝒯 obtained from the real data, provided in the TARO R package. We constructed four different settings; in each setting, the true signal is sparse, with only 5% of features affecting the multivariate outcome, but the set of important features differs in its properties. Specifically, we simulate the unit-rank components of the coefficient matrix such that the following feature sets are relevant: a) features with higher variation, b) rare features, c) fine-resolution features (leaf nodes), and d) aggregated features (internal nodes). We set the sample size *n* to 300. The number of microbial genera *p* varied in the range from 200 to 225 depending on the construction of **C**. Finally, we generated the response matrix **Y** using the true model (1) with error term **E** simulated at a signal-to-noise ratio (SNR) of 0.5, following the definition of the SNR in Mishra et al. [2017].

To provide insight into the relative performance of TARO, we consider several alternative procedures:

**TRAC**: tree-aggregation of compositional data [Yan and Bien, 2021], which is designed for a single outcome.

**CRRR**: linear-constrained reduced-rank regression, a simplified version of TARO without feature selection.

**SeCURE**: sequential co-sparse factor regression [Mishra et al., 2017], which is not designed for the microbiome setting.

For an overview of the method properties, see Table 1. Comparing TARO with the marginal approach of TRAC emphasizes the relevance of joint modeling of the multivariate outcome, while the comparison to CRRR highlights the significance of the sparsity-inducing penalty when compared with TARO. Finally, the comparison to SeCURE showcases the importance of imposing linear constraints due to compositionality.

**Table 1.** Summary of models compared.

|  | Multivariate model | Outcome selection | Tree-guided feature selection | Microbiome data as compositional |
| --- | --- | --- | --- | --- |
| **TARO** | ✓ | ✓ | ✓ | ✓ |
| TRAC | ✗ | ✗ | ✓ | ✓ |
| CRRR | ✓ | ✗ | ✗ | ✓ |
| SeCURE | ✓ | ✓ | ✗ | ✗ |

We compare the model results in terms of error in estimating the coefficients 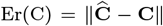, prediction error 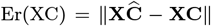, and feature selection. Performance in feature selection is based on comparing the sparsity pattern of 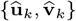 to *{***u**_*k*_, **v**_*k*_*}* in terms of the false positive rate (FPR) and false negative rate (FNR).

Across fifty replicates of setting a), where features with higher variation are associated with the multivariate outcomes, TARO achieves consistently lower estimation and prediction error than alternative methods (Figure 2). TARO also achieves a reasonable balance betwen the false negative rate (FNR) and false positive rate (FPR) for feature seleection (Table 2). Results from other simulation settings are provided in Figure S1 of the Supplementary Material.

**Table 2.** Performance comparison in terms of coefficient estimation error Er(C), prediction error Er(XC), and the false positive rate (FPR) and false negative rate (FNR) for feature selection.

| Model | $\text{Er}(\mathbf{C})$ | $\text{Er}(\mathbf{XC})$ | FNR | FPR |
| --- | --- | --- | --- | --- |
| **TARO** | **1.3** | **240** | **0.180** | **0.068** |
| TRAC | 9.1 | 650 | 0.031 | 0.670 |
| CRRR | 410.0 | 1800 | 0.000 | 1.00 |
| SeCURE | 5.2 | 1400 | 0.890 | 0.010 |

**Fig. 2:**
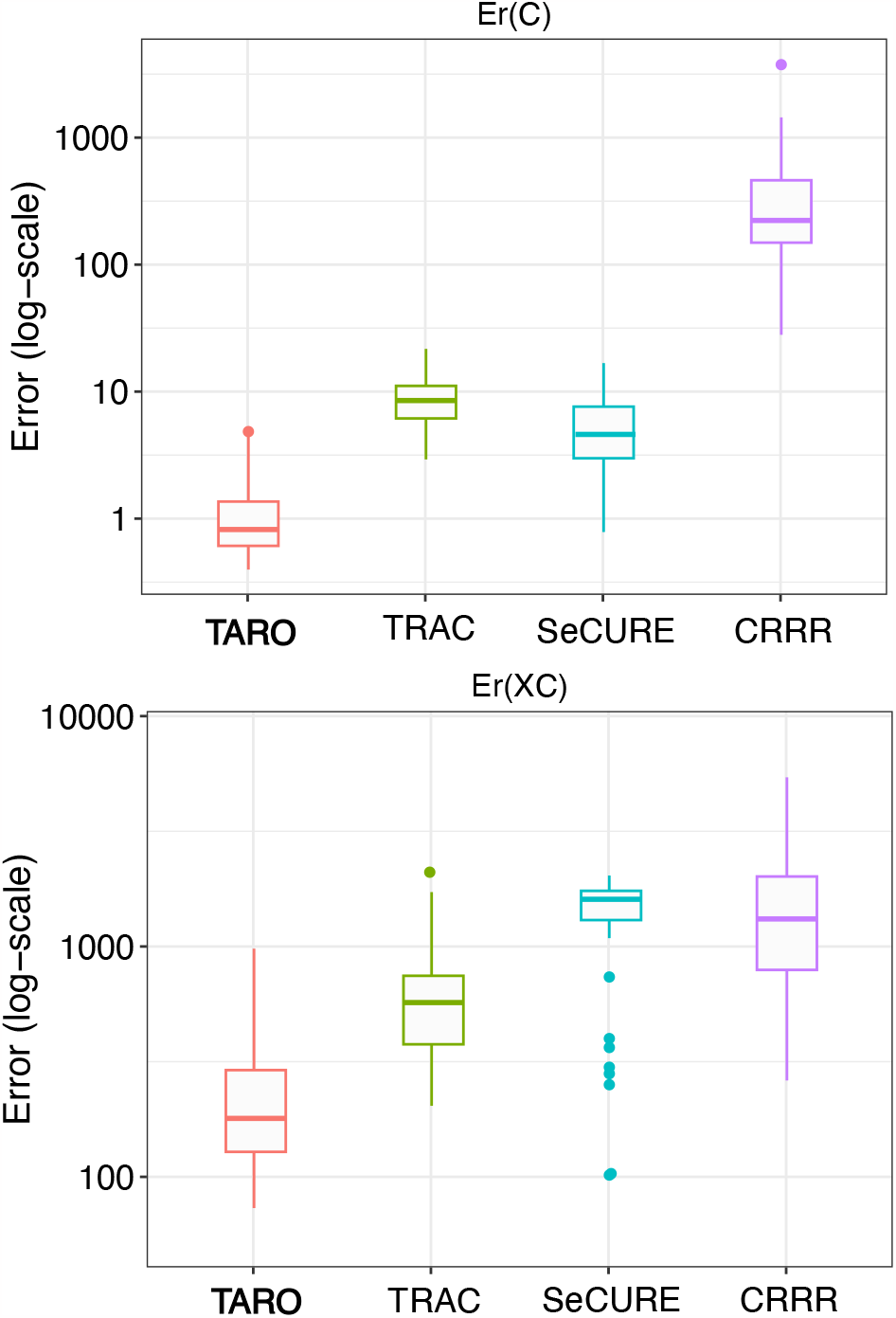
Boxplots of the coefficient estimation error Er(C) and prediction error Er(XC).

Compared to existing approaches for modeling the multivariate outcome with compositional covariates as predictors, TARO demonstrates superior performance in estimation error, prediction error, and sparsity recovery. In high-dimensional settings where the underlying association can be expressed in terms of a low-rank and sparse coefficient matrix, the superior performance of TARO to CRRR and TRAC shows the usefulness of joint modeling of multivariate outcomes and the sparsity-inducing penalty.

### Analysis of colorectal cancer data using TARO

There is increasing evidence that the human gut microbiome influences diseases including colorectal cancer and inflammatory bowel disease through the production of metabolites [Lee-Sarwar et al., 2020]. To provide insight into microbial-metabolite relationships in the gut ecosystem, we applied TARO to analyze metagenomic and metabolomic profiling data collected from participants undergoing colonoscopy as part of a large-scale study in colorectal cancer [Yachida et al., 2019]. A processed version of this data is provided through the curated gut microbiome-metabolome data resource https://github.com/borenstein-lab/microbiome-metabolome-curated-data/ [Muller et al., 2022]. The processed data includes observations for *n* = 347 participants on *q* = 249 metabolites and *p* = 1456 microbial genera. We defined the adjacency matrix **A** using the taxonomic tree relating the observed genus-level features.

To provide new insight into this complex data set, we applied TARO to characterize the interplay between microbiome profiles and metabolites. As a contrast to TARO, which performs a joint analysis to identify latent factors, we applied sparse principal component analysis (PCA) [Erichson et al., 2020] separately on the **X** and **Y** matrices. Our goal was to find sets of microbiome features and metabolites that work together to impact phenotypes of interest. We computed the top eight principal components from independent PCA and aligned the components based on their correlation (Figure 3A). However, the resulting cloud of points suggests that the independently inferred latent components may be capturing distinct activity within each modality, rather than shared processes.

**Fig. 3:**
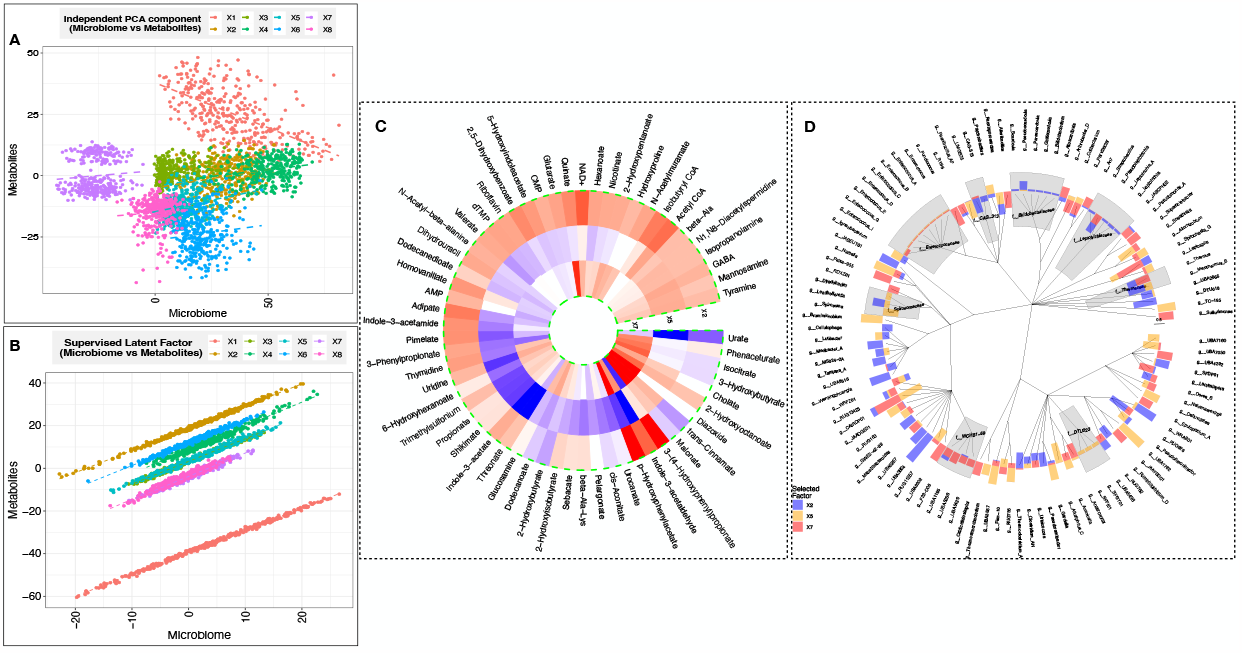
(A) Scores for each sample from PCA conducted independently on each modality. (B) Scores for each sample from TARO model. (C) Heatmap of coefficients of selected metabolites for clinically relevant factors. (D) Circular barplot of coefficients of selected microbiome features for clinically relevant factors.

Using TARO, we were able to identify association patterns represented by a low-rank and sparse estimate of the coefficient matrix. Each unit-rank component within the coefficient matrix provides valuable information regarding the subset of metabolites (via the sparse estimate of the loading matrix **V**) that directly correspond to a subset of microbiome features (via the sparse estimate of **U**). The latent factors identified by TARO capture microbiome-metabolite relationships more efficiently than those from independent PCA, as each latent factor represents a set of microbiome and metabolite features working in concert (Figure 3B).

Each of the eight latent factors identified by TARO represents distinct sets of microbes and metabolites working together to perform specific tasks in the gut. To identify the latent factors with the greatest clinical relevance, we fit a logistic regression model with the scores on the latent factors as predictors and sample classification (colorectal cancer vs. normal tissue) as the response. We adjusted for age, sex, and body mass index (BMI) in the model. Three latent factors (X2, X5, and X7) were significant (*p <* 0.05) in the logistic regression model (Figure S2 of the Supplementary Material). TARO selects a sparse set of metabolites (Figure 3C) and microbiome features (Figure Figure 3D) that contribute to these clinically relevant latent factors. TARO identifies both both genus-level and also aggregated features are identified as important (Figure 3D).

We next sought to characterize possible biological processes represented by these latent factors. Based on metabolite set enrichment analysis (Figure S3 of the Supplementary Material), we identified propanoate metabolism as a key metabolic pathway represented by X2. Propionate is an abundant short chain fatty acid in the gut, and altered propionate metabolism has been linked to cancer progression and aggressiveness [Gomes et al., 2022]. The family Bifidobacteriaceae, which includes the genus *Bifidobacterium*, was identified as an aggregated feature that contributes to X2 (Figure 3D). Although Bifidobacteriaceae do not directly produce propionate, they contribute to propionate and butyrate production through cross-feeding in the gut [Scott et al., 2014]. Enrichment analysis revealed that X5 captures functions related to energy metabolism including the citric acid cycle and beta oxidation of very long chain fatty acids. The genus UBA1762 was identified as a microbial feature contributing to X5; UBA1762 belongs to the family Ruminococcaceae, which has been previously identified as a taxonomic feature positively associated with response to cancer immunotherapy [Gopalakrishnan et al., 2018]. Finally, pyrimidine metabolism, which has been closely linked with cancer progression [Wang et al., 2021], was identified as a key pathway in X7. Interestingly, several cancer drugs, including the chemotherapeutic agent fluoropyrimidine, act to disrupt pyrimidine metabolism [Spanogiannopoulos et al., 2022]. The family WCHB1.69, which belongs to the order Bacteroidales, is identified as an important microbial feature for X7; Bacteroidales play an important role in shaping response to immmunotherapy [Vétizou et al., 2015].

In summary, TARO enables us to identify a small number of latent factors that are relevant to colorectal cancer status and the specific microbiome and metabolite features represented in each factor. TARO enables the formulation of testable hypotheses regarding the interplay between the microbiome and metabolome. The TARO results highlight potential avenues of intervention that can be further explored through pre-clinical studies in mice.

## Conclusion

TARO provides an effective tool for the integration of microbiome and metabolite data sets. Through a specially designed penalization approach, TARO is able to identify specific features from each modality that contribute to a small set of latent factors. Importantly, TARO respects unique aspects of microbiome data including its compositionality and the tree-structured relationships among features. We illustrate the superior performance of TARO in simulation settings and discuss its application to a colorectal cancer data set.

More broadly, TARO may be applied for the integration of microbiome profiles with high-dimensional data types other than metabolomics. For example, **Y** could instead represent microbial functional proteins (metaproteomics) or host-associated factors such as immune cell abundances. An interesting possible extenion of TARO would be to acknowledge structure among the **Y** variables, such as pathway membership or network relations, in addition to structure on the **X**.

## Supplementary data

Additional details of the TARO algorithm are provided in the Supplementary data file.

## Data availability

The metabolite and microbiome profiles analyzed in the case study were originally described in Yachida et al. [2019]. A processed version of this data set has been shared through the curated gut microbiome-metabolome data resource [Muller et al., 2022] and is available online at https://github.com/borenstein-lab/microbiome-metabolome-curated-data/wiki/.

## Supplementary

### Weighted adaptive-elastic-net penalty

We use the adaptive elastic net penalty [**??**],

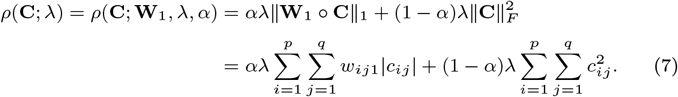

Here ‖ ·‖_1_ denotes the *ℓ*_1_ norm, the operator “◯” stands for the Hadamard product, **W**_1_ = [*w*_*ij*1_]_*p×q*_ is a pre-specified weighting matrix, *λ* is a tuning parameter controlling the overall amount of regularization, and *α ∈* (0, 1) controls the relative weights between the two penalty terms. We set 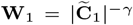 such that 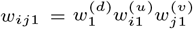, with

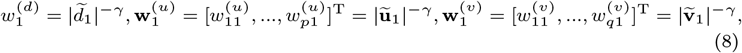

where 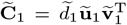 is the first set of unit-rank RRR estimators and *γ* is a non-negative constant with | · |^*−γ*^ componentwisely defined.

### URE-TARO procedure

Here, we provide additional detail on the unit rank estimation procedure for TARO described in Section 2.2 of the main manuscript. In our previous work on sparse factor regression Mishra et al. [2021], we imposed sparsity directly on **C**. To enable appropriate aggregation of the microbiome features in the factor regression framework, we instead propose to impose sparsity on **Γ**.

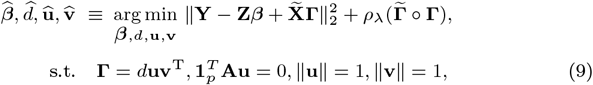

In terms of the parameters *{****β***, *d*, **u, v***}*, the optimization problems of URE-TARO is a multi-convex problem. We define the weighted penalty as:

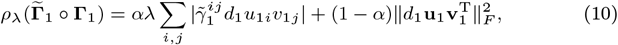

where *λ* is the tuning parameter, *α* provides a relative weights of *ℓ*_1_ and *ℓ*_2_ penalty. We estimate the model parameters using an iterative procedure that cycles between **u**-step, **v**-step and ***β***-step until convergence.

#### u-step

For the fixed **v** and ***β***, we jointly update (*d*, **u**) satisfying ‖**v**‖ = 1. Let us define 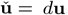. In terms of 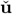, the optimization problem (9) is equivalent to solving (constrained adaptive elastic-net problem)

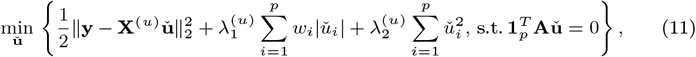

where **y** = vec(**Y** *−* **Z*β***), **X**^(*u*)^ = **v** ⊗ **X**, 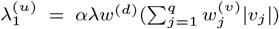, and 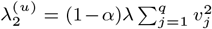. Here vec(·) is the vectorization operator, and ⊗ denotes the Kronecker product. We recover the singular value estimate as 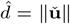 and singular vector estimate as 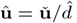.

#### v-step

For fixed **u** and ***β***, we minimize the objective function in terms of the block variable (*d*, **v**) such that ‖**u**‖ = 1. Let us define 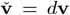. In terms of 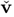, the optimization problem (9) is equivalent to solving (an adaptive elastic-net problem)

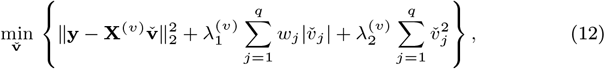

where **X**^(*v*)^ = **I**_*q*_ ⊗ (**Xu**), 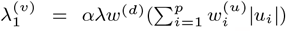, and 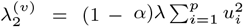. We recover the singular value estimate as 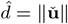 and singular vector estimate as 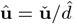.

#### *β*-step

For fixed {*d*, **u, v**}, unique solution minimizing the objective function is given by

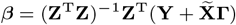

### Constrained adaptive elastic-net solution

Consider a genetic form of the linear constrained adaptive elastic net regression,

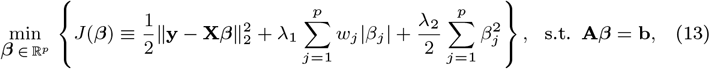

where **y** *∈* ℝ^*n*^, **X** *∈* ℝ^*n×p*^, **A** = (**a**_1_, …, **a**_*p*_) *∈* ℝ^*h×p*^, **b** *∈* ℝ^*h×*1^, **w** = [*w*_1_, …, *w*_*p*_]^T^ are some predetermined weights, and *λ*_1_ and *λ*_2_ are tuning parameters.

This is an equality-constrained convex optimization problem. The augmented Lagrangian function is

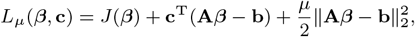

where **c** is the Lagrange multiplier, and *μ >* 0 is a penalty parameter. The iterative steps to solve the problem,

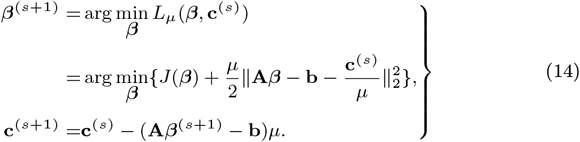

The method converges under very general conditions. As the iteration proceeds, the residual **A*β***^*s*+1^ *− b* converges to zero, yielding optimality.

Following (14), the key is to minimize

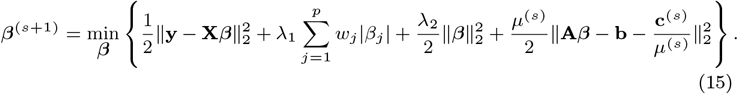

Here the penalty parameter *μ*^(*s*)^ can be updated along the interactions; let *μ* → ∞ or increase with small increments can in general improve the speed of convergence [**?**]. The above problem can be efficiently minimized by a coordinate descent algorithm. Suppose all the *β*_*k*_s are fixed except *β*_*j*_, and denote 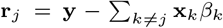. The objective function with respect to *β*_*j*_ becomes

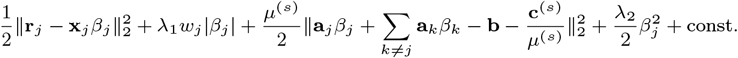

Then it can be easily verified that

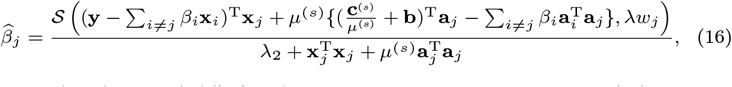

where 𝒮 (*m, λ*) = sign(*m*)(|*m*| *− λ*)_+_ is the soft-thresholding operator. (15) can then be solved by iteratively updating each *β*_*j*_, *j* = 1, …, *p*, by (16) until convergence. Our proposed algorithm is presented in Algorithm 0.

#### Algorithm 2 Bregman Coordinate Descent Algorithm (BCDA)

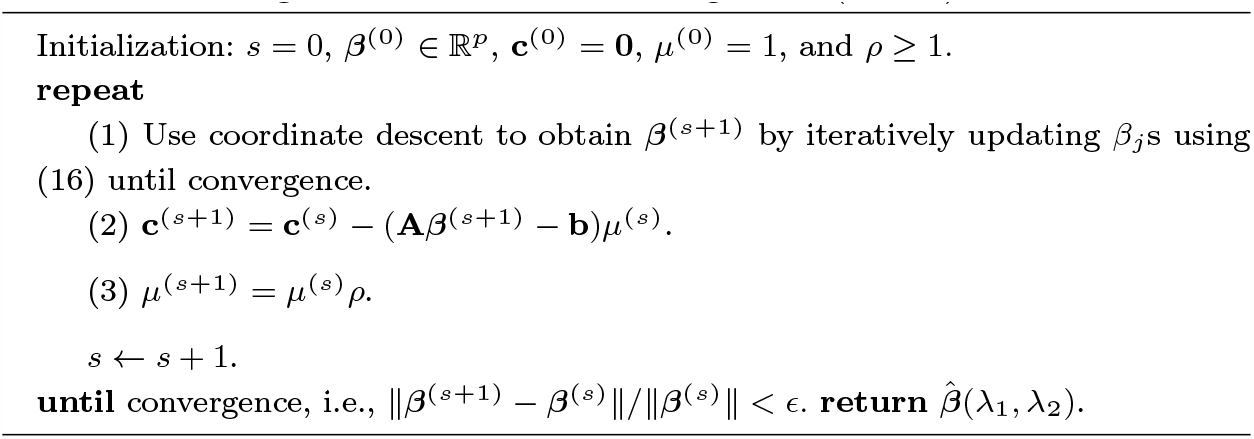

### Constrained reduced-rank regression

We consider a general form of the linear-constrained reduced rank regression as

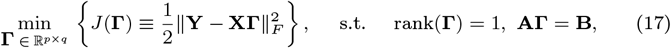

where **Y** *∈* ℝ^*n×q*^, **X** *∈* ℝ^*n×p*^, **A** = (**a**_1_, …, **a**_*p*_) *∈* ℝ^*h×p*^ and **B** *∈* ℝ^*h×q*^. To solve the optimization problem, we write the augmented Lagrangian function as

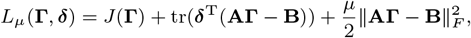

where ***δ*** *∈* ℝ^*h×p*^ is the Lagrange multiplier, and *μ >* 0 is a penalty parameter. The iterative steps to solve the problem,

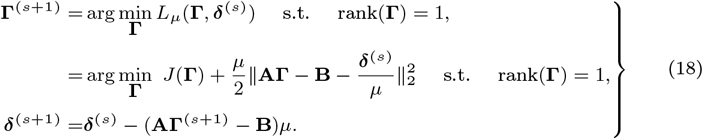

The method converges under very general conditions. As the iteration proceeds, the residual *‖***AΓ**^*s*+1^ *−* **B***‖* converges to zero, yielding optimality. We simplify the rank constrained *L*_*μ*_(**Γ, *δ***^(*s*)^) as

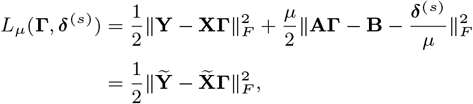

where 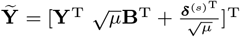 and 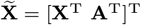. An optimal solution of

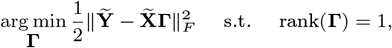

is given by 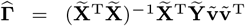 where 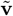 is the largest eigen-vector of 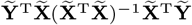. Our proposed algorithm is presented in Algorithm 3.

#### Algorithm 3 Linear Constrained Reduced Rank Regression

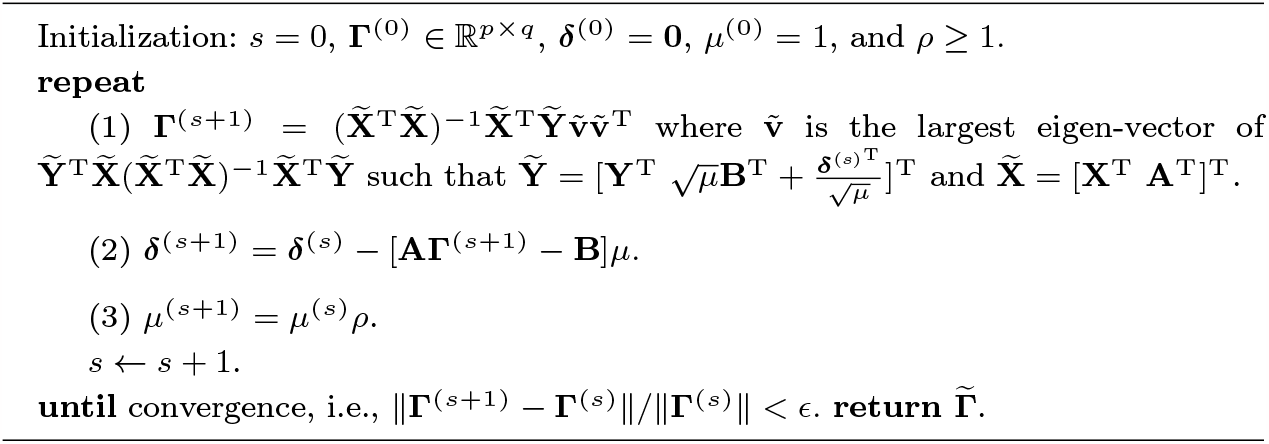

## Supplementary figures

### Simulation results

Here, we provide additional figures summarizing the simulation results for the settings described in Section 3.1 of the main manuscript. Specifically, we considered simulation settings where the unit-rank components of the coefficient matrix are constructed such that the following feature sets are relevant: a) features with higher variation, b) rare features, c) fine-resolution features (leaf nodes), and d) aggregated features (internal nodes). The results for setting a) are given in Figure 2 of the main manuscript. The results for the remaining scenarios are provided in Figure 4. TARO consistently outperforms the alternative methods considered in terms of accuracy in the estimation of the true coefficient matrix, prediction accuracy, and recovery of the true features.

**Fig. 4:**
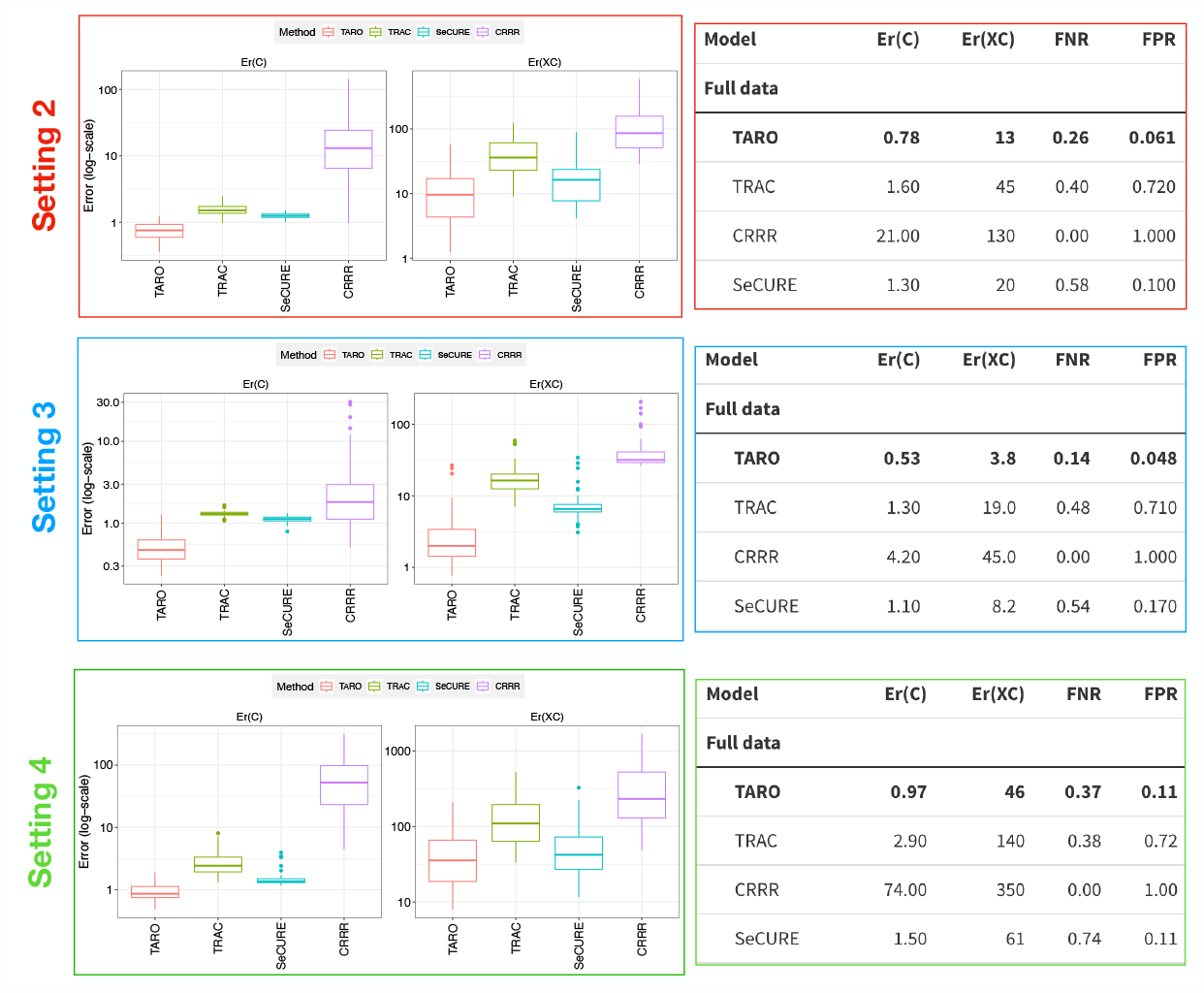
Model performance comparison in various simulated settings in terms of estimation accuracy, prediction, and sparsity recovery.

### Application results

We provide a snapshot of the logistic regression modeling results in the upper left of Figure 5. Rectangular heatmaps showing the contribution of input features to the clinically relevant latent factors are shown for the metabolite variables (upper right) and microbiome features (bottom). The results of metabolite set enrichment analysis are shown in Figure 6.

**Fig. 5:**
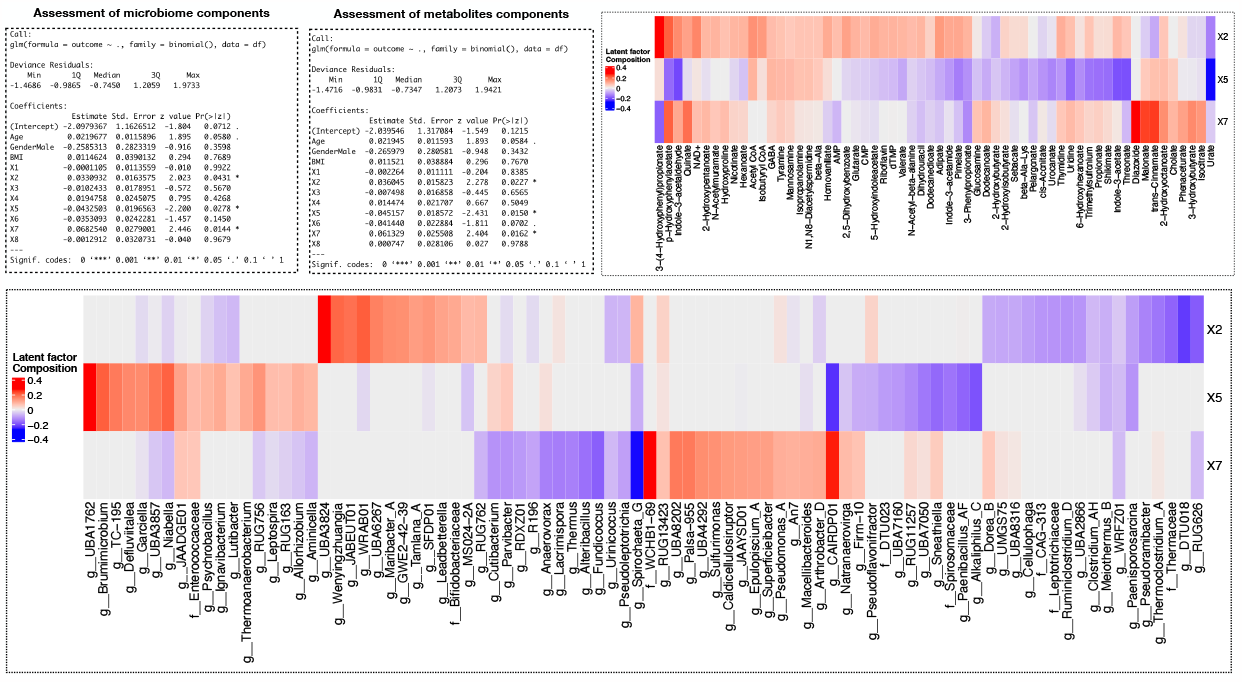
Assessment of the eight microbiome/metabolites latent factors in association with the outcome of interest (healthy vs colorectal cancer patients). Heatmaps show the selected microbiome and metabolites in respective latent factors.

**Fig. 6:**
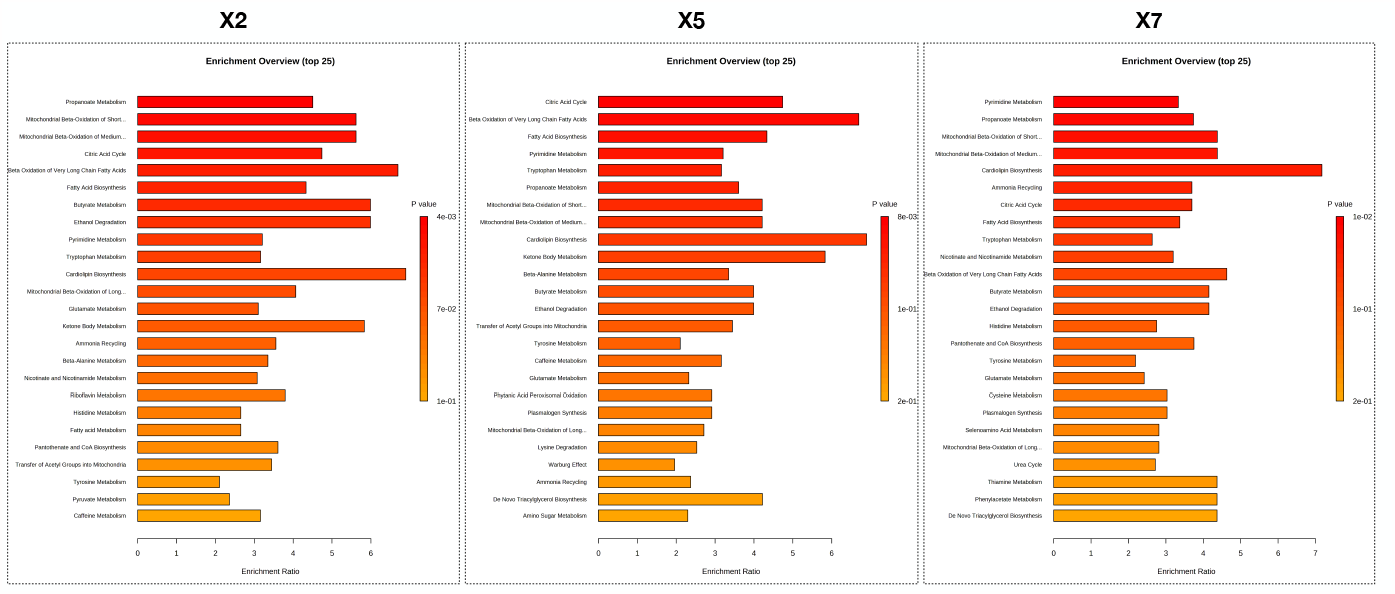
Metabolic pathway obtained using metabolite set enrichment analysis.

